# Lipid Nanoparticles with Aptamers Enable Targeted mRNA Delivery to CD4^+^ T Cells

**DOI:** 10.1101/2025.09.10.675359

**Authors:** Shrey Shah, Meenakshi Ranasinghe, John Decker, Keith Fraser, Ying Wang, Sarah Kinlaw, Morgan Barker, Sawan Patel, Adam Friedman, Sherwood Yao

## Abstract

*In vivo* genetic engineering of T cells could overcome the logistical, biological, and safety challenges of *ex vivo* modification, but effective and safe delivery systems remain limited by a lack of cellular specificity. Here, we developed aptamer-functionalized lipid nanoparticles (LNPs) for targeted mRNA delivery to CD4+ T cells, employing both a validated CD4-binding aptamer (Apt62) and novel aptamers generated using our proprietary transformer-based AI language model, AptaBLE. LNPs formulated with ionizable lipid SM102 or MC3 and conjugated with aptamers at controlled densities were physiochemically characterized and assessed for binding, in vitro transfection, in vivo biodistribution, and safety evaluation. Aptamer-functionalized LNPs demonstrated selective nanomolar binding to recombinant CD4, achieved enhanced transfection of CD4^+^ versus CD4^-^ T cells in vitro, and significantly enriched mRNA delivery to immune-rich tissues in vivo, achieving up to 70-fold spleen signal enhancement with SM102 formulations compared to non-targeted controls, while maintaining suitable safety profiles. Overall, these findings demonstrate aptamer-functionalized LNPs, augmented by AI-guided aptamer design, as a tunable, non-immunogenic platform for in vivo T cell engineering.

**Highlights:**

- Aptamer-functionalized LNPs enable selective mRNA delivery to CD4^+^ immune cells rich organs.
- Aptamer-functionalized LNPs maintain a favorable systemic safety profile *in vivo*.
- AI-guided AptaBLE platform generated functional aptamers validated in nanoparticle delivery.

## 1. Introduction

Genetic modification of T cells has enabled transformative immunotherapies, regulating immune responses to treat hematological malignancies, autoimmune disorders, and infectious diseases.^1–3^ Among various T cell subsets, CD4^+^ T cells play a central role in orchestrating immune responses by activating cytotoxic T cells and optimizing their responses, enhancing B cell function, and maintaining immune homeostasis.^4^ Despite their critical role, the most commonly used approach of genetically modifying T-cells *ex vivo* presents significant challenges.^5^ T cells must first be isolated from a heterogeneous pool of peripheral blood mononuclear cells, followed by selection of CD4^+^ or CD8^+^ subsets using antibody-based sorting techniques. ^6–8^ The purified cells then undergo weeks of proliferation under controlled conditions, genetic modification via viral or non-viral vectors, and eventual reinfusion into patients. This process is not only labor-intensive and costly but also introduces biological challenges, including heterogeneity in modified cell phenotypes, variable survival, and loss of sustained function post-infusion.^6–8^ Moreover, reinjection of genetically modified T-cells can trigger excessive cytokine release, leading to severe toxicities such as cytokine release syndrome and immune effector cell-associated neurotoxicity syndrome.^9,10^ These inflammatory responses driven by interleukin-6 (IL-6), interferon-gamma (IFN-γ), and tumor necrosis factor-alpha (TNF-α) can result in systemic immune dysregulation, vascular leakage, and in severe cases, multi-organ dysfunction.^11–13^ The risk associated with T-cells reinfusion, along with logistical and biological constraints of *ex vivo* T-cell engineering, underscores the need for alternative strategies that enable direct *in vivo* genetic programming of T cells while preserving their physiological state and function.

Lipid nanoparticles (LNPs) have emerged as a promising non-viral gene delivery platform, with their clinical success in mRNA vaccines establishing their potential for broader therapeutic applications.^14,15^ Their biocompatibility, tunable characteristics, and the ability to encapsulate and deliver mRNA make them a viable option for gene therapy, particularly for T-cell modulation.^14,16^ However, conventional LNPs exhibit high protein expression in the liver, resulting in low expression and targeting to CD4^+^ T cell-rich organs such as the spleen, bone marrow, and thymus.^17^ To overcome these limitations, endogenous targeting strategies such as Selective Organ Targeting (SORT) have been developed by incorporating supplemental lipids into LNP formulations to modify LNP surface and tune the protein corona composition to deliver mRNA towards non-hepatic tissues.^18^ While SORT enhances organ-level tropism, its ability to achieve cellular specificity, particularly for immune cell subsets, remains limited.^19^ This has led to growing interest in active targeting approaches, wherein LNPs are surface-decorated with ligands that enable receptor-specific uptake by desired cell types. Among these, monoclonal antibodies (mAbs) have been widely explored to direct LNPs toward immune cells.^20–22^ However, mAb functionalized LNPs have several fundamental drawbacks, including Fc-mediated immune activation, steric hindrance affecting LNP uptake, and high manufacturing costs.^23–25^ For example, It has been seen that the pharmacokinetic and pharmacodynamic profiles of antibodies are altered when they are functionalized to nanoparticles.^26^ The first administration of antibody functionalized nanoparticles shows prolonged circulation, but with repeated dosing, immune responses are triggered that drive progressively faster clearance.^26^ In contrast, the free antibody maintains favorable pharmacokinetics and does not induce such immunogenicity.^26^ Peptides have also been investigated as targeting ligands, offering smaller size and ease of synthesis, but their utility *in vivo* is often limited by proteolytic degradation, relatively short circulation time, modest receptor affinity, and peptide misfolding and aggregation.^27–29^

Recognizing the limitations of antibodies and peptides, there is increasing interest in alternative ligands that combine high specificity and ease of synthesis. Aptamers, short single stranded oligonucleotides that fold into defined three dimensional structures, satisfy these criteria.^30,31^ Their small size, low immunogenicity, and chemical stability have enabled broad use in diagnostics and biosensing, where they have already been applied successfully.^32,33^ Recent advances have extended the utility of aptamers from diagnostics into therapeutic applications, particularly in oncology and immuno-oncology.^34^ RNA aptamers directed against epidermal growth factor receptor have been shown to recognize clinically relevant mutant variants such as L858R and T790M and to enable selective tumor targeting in non–small cell lung cancer models.^35^ Nucleolin binding DNA aptamers such as AS1411 have progressed into phase I and II clinical trials, where they demonstrated favorable safety and preliminary antitumor activity.^36^ Although these studies demonstrate that aptamers can engage to extracellular receptors with high affinity and promote cellular uptake, they have been under explored as targeting ligands. Thus, in this study we evaluated whether functionalizing lipid nanoparticles with aptamers could enable selective delivery of mRNA therapeutics to defined immune cell subsets.

Rather than immediately developing a de novo sequence, we adopted a proof-of-concept strategy using a well-validated human CD4-binding aptamer (Apt62).^37^ This aptamer was originally identified by SELEX and next-generation sequencing and has been shown to bind CD4 with nanomolar affinity, disrupt gp120– CD4 interactions, and inhibit HIV-1 entry across multiple CD4-expressing cell lines.^38^ Importantly, it also demonstrated stability in human serum, underscoring its suitability for systemic administration. Building on this prior validation, we repurposed Apt62 as a benchmark ligand to evaluate whether aptamer-mediated surface modification of LNPs could improve targeted mRNA delivery to CD4^+^ cells and immune-rich tissues such as the spleen following systemic administration. Although in this study we chose to use a previously validated aptamer, a major limitation in aptamer discovery lies in the PCR-driven Systemic Evaluation of Ligands by Exponential Enrichment SELEX process.^39,40^ It is time-consuming, typically requiring 1–3 months of iterative rounds, and the probability of isolating high-affinity binders is often low.^39,40^ To improve discovery throughput and expand the sequence design space, we further applied AptaBLE, a sequence-first machine learning framework trained on diverse aptamer protein datasets, which can generate new CD4 binding DNA aptamers within weeks.^41^ These computationally designed candidates were then evaluated alongside Apt62 for their capacity to drive CD4-selective binding and mRNA delivery.

### Materials

Ionizable lipids SM-102 (Cat. No. BP-25499) and MC3 (Cat. No. BP-25497), DSPC (Cat. No. BP-25623) were purchased from Broad Pharm (San Diego, CA). Cholesterol (Cat. No. 700100) and PEGylated lipid (PEG2000-DMG, Cat. No. 880151) were obtained from Avanti Polar Lipids (Alabaster, AL). DSPE-PEG2000-Maleimide (Cat. No. 2049) was obtained from Nanosoft Polymers (Winston-Salem, NC). GFP-encoding mRNA (Cat. No. RP-A00009) and firefly luciferase (fLuc) mRNA (Cat. No. RP-A00023) were purchased from GenScript (Piscataway, NJ). All aptamers and molecular beacons were synthesized and purchased from Integrated DNA Technologies (Morrisville, NC). Tris(2-carboxyethyl) phosphine (TCEP-HCL) was purchased from Gold Biotechnology (Olivette, MO) and N-Succinimidyl-S-acetylthioacetate (SATA, Cat. No. A508892) was purchased from Ambeed (Buffalo grove, IL).

SUPT-1 (CD4^+^) and HSB-2 (CD4^-^) human cell lines, Roswell Park Memorial Institute medium (RPMI - 1640), and Iscove’s Modified Dulbecco’s Medium (IMDM) culture media were obtained from the American Type Culture Collection (Virginia, USA). Fetal Bovine Serum (FBS), the LIVE/DEAD™ Fixable Far Red Dead Cell Stain Kit, anti-human CD3-FITC monoclonal antibody, Human CD4 monoclonal antibody (Cat. No. 69-0049-42), mouse CD4 monoclonal antibody (Cat. No. 14-0042-82), d-Luciferin, and Antibiotic-Antimycotic were purchased from ThermoFisher Scientific (Waltham, MA). The CellTiter-Glo® Luminescent Cell Viability Assay was obtained from Promega (Madison, WI). Female C57BL/6 mice, aged 6 to 8 weeks, were obtained from Charles River Laboratories (Wilmington, MA) for *in vivo* experiments. ELISA kits for the quantification of IL-6, TNF-α, and alanine aminotransferase (ALT) were purchased from Abcam. Ethanol, Phosphate buffer saline (PBS, 20X), Tris-EDTA buffer (TE, 20X), HEPES buffer (1M), NaCl, MgCl2, were purchased from VWR (Radnor, PA).

## 2. Methods

### 3.1 Synthesis of Non-targeting Lipid Nanoparticles (LNPs)

mRNA-encapsulated lipid nanoparticles were formulated using a previously published method with slight modifications.^22,42^ Briefly, the ethanol phase was prepared by combining ionizable lipid (SM102 or MC3), DSPC, cholesterol, DMG-PEG2000, and DSPE-PEG2000-MAL in 50:10:38.5:1.2:0.3 molar ratio and the aqueous phase mRNA (N/P= 10) in citrate buffer (5 mM, pH 4.0). LNPs were formed by mixing an ethanolic lipid solution with an aqueous mRNA solution using a microfluidic device (Pump 33 DDS, Harvard Apparatus, MA) at a 1:3 ratio, followed by dilution in citrate buffer (5 mM, pH 4.0). The formulation was then dialyzed against 1× PBS (pH 7.5) to remove ethanol and confirm buffer exchange. Post-dialysis LNPs were concentrated via centrifugation at 1000 × g using 100 kDa Vivaspin filters (Sartorius, Germany) prior to aptamer conjugation.

### 3.2 Synthesis of Aptamer-Conjugated LNPs

Aptamer conjugation to lipid nanoparticles (LNPs) was performed via thiol–maleimide chemistry. Thiol-modified aptamers in aptamer folding buffer (50 mM HEPES, 5 mM MgCl_2_, 150 mM NaCl, pH 7.4) were folded using a heat-annealing method consisting of heating to 95 °C, rapid cooling to 4 °C, and gradual heating to 25 °C. Disulfide bonds on the folded aptamers were reduced using reducing agent tris(2-carboxyethyl) phosphine (TCEP) at a 1:10 molar ratio of aptamer to TCEP. Unreacted aptamers were purified by dilution in 1× PBS (pH 7.5) followed by centrifugation, as described previously.^43^ The aptamer– LNP formulations were prepared at theoretical aptamer-to-LNP ratios of 25, 75, and 100, denoted at the end of each formulation name (e.g., SM102-Apt62:75). The sequences of the aptamers used for conjugation are provided in Table S1 of the Supporting Information.

### 3.3 Synthesis of Antibody-Conjugated LNPs

Antibody conjugation to LNPs was performed using SATA modification followed by thiol–maleimide chemistry.^22^ Briefly, antibodies were modified with N-succinimidyl S-acetylthioacetate (SATA) at a 1:10 antibody-to-SATA molar ratio in 1× PBS buffer (pH 7.5) and incubated it 45 minutes to introduce protected sulfhydryl groups. Deprotection was carried out using hydroxylamine buffer (0.5 M hydroxylamine, 25 mM EDTA in 1× PBS, pH 7.5) by adding 1 volume of hydroxylamine buffer to every 10 volumes of antibody solution and incubating for 2 hours, thereby exposing free thiols. Excess reagents were removed using spin desalting columns (Cat. No. 89882, ThermoFisher Scientific, USA). The resulting sulfhydryl-functionalized antibodies were conjugated to maleimide groups on the LNP surface through thioether linkage. Final purification of antibody-conjugated LNPs was performed either by centrifugation at 1000 × g using 300 kDa Vivaspin centrifugal filters (Sartorius, Germany) or by size-exclusion chromatography using Sepharose CL-4B gel filtration columns (G-Biosciences, USA).

### 3.4 Physicochemical characterization of LNPs

LNPs were then analyzed using dynamic light scattering (DLS) performed on a Zetasizer Nano (Malvern Instruments, Malvern, UK) to determine their diameter (z-average), polydispersity index (PDI), and zeta potential. The mRNA concentration and encapsulation efficiency of the LNPs were measured using Quant-iT RiboGreen RNA assay (Cat. No. R11491, ThermoFisher Scientific, USA). Aptamer density on LNP was measured using an in-house developed molecular beacon method.

Briefly, a molecular beacon complementary to a unique ∼12-nucleotide loop region of the aptamer was designed, incorporating a 5′-fluorescein (FAM) fluorophore and a 3′-Black Hole Quencher 2 (BHQ2). The beacon was synthesized and HPLC-purified (Integrated DNA Technologies, USA) before use in the following assay. Serial dilutions of aptamer (0–800 nM) and molecular beacon (1 µM) were prepared in aptamer folding buffer (50 mM HEPES, 5 mM MgCl_2_, 150 mM NaCl, pH 7.4). Aptamer or aptamer–LNP samples (14 µL) were mixed with molecular beacon solution (14 µL) and Triton X-100 (3 µL, 0.5%) in a clear PCR 96-well plate. Fluorescence was measured using a qPCR instrument (QuantStudio3, Applied Biosystems by ThermoFisher Scientific, USA) under kinetic mode with a thermal profile of 90 °C (denaturation) to 30 °C (cooling) at 1.6 °C/s. Aptamer density on LNPs was determined from a calibration curve generated with free aptamer standards.

Antibody conjugation to LNP was confirmed using a fluorescence-labeled antibody (Human CD4 Monoclonal Antibody (RPA-T4) eFluor 506, Thermo Fisher Scientific, Cat No.69-0049-42).

### 3.5 Biolayer interferometry (BLI)

BLI was used for the kinetic characterization of Apt62 and its MC3-based lipid nanoparticle (LNP) formulations using a Gator Bio® Prime 8-channel instrument (Gator Bio, Palo Alto, CA). All assays were performed at 30 °C in 96-well BLI 96-Flat Black Plates, with samples prepared in assay buffer containing 50 mM HEPES, 150 mM NaCl, and 5 mM MgCl_2_ (pH 7.4). For kinetic analysis of free Apt62, anti-VHH biosensors were used to immobilize anti-human CD4 nanobodies (sequence: EVQLVESGGGLVQPGGSLRLSCAASGFTFSKLAMSWHREPPGKGREWLADIDSSGDTTDYLASVKGRFTISRDNAKNTLYLQMDSLKSEDTGVYYCASREDPPGYWGQGTQVTVSS; expressed and purified by Bon Opus Biosciences, LLC) at 50 µg/mL for 180 seconds, followed by capture of 6x-His-tagged recombinant human CD4 (rCD4; Sino Biological, Inc.) at 3 µg/mL for 180 seconds. This immobilization strategy was employed to avoid steric hindrance arising from the proximity of the Aptamer 62-binding epitope to the 6xHis tag on CD4, which interfered with direct binding when CD4 was immobilized via the His tag. Indirect immobilization via the nanobody allowed for a more ideal orientation of the target protein and accurate kinetic assessment. Aptamer 62 was then tested at concentrations of 20,000, 10,000, 5000, 2500, 1250, and 625 nM, with association and dissociation phases of 900 seconds each.

For kinetic analysis of MC3-based LNPs surface modified with Apt62, anti-His biosensors were used to immobilize 6x-His-tagged rCD4 (3 µg/mL) for 180 seconds, followed by exposure to aptamer-functionalized LNPs at aptamer-equivalent concentrations of 364, 182, 91, 45.5, 23, and 11 nM, with 600 seconds for both association and dissociation. Kinetic analysis of Aptamer 62-immobilized MC3-based LNPs was possible because of avidity effects; LNPs bearing ∼20 to >100 Aptamer 62 ligands were able to interact strongly with the cognate epitopes on 6xHis-hCD4 in aggregate despite the aforementioned steric hindrances that abrogate the observation of 1:1 Aptamer 62-6xHis hCD4 binding kinetics. All samples were run in column format, and data were acquired using GatorOne software. Reference sensor signals (assay buffer only) were subtracted, and the resulting curves were processed using Savitzky-Golay filtering and fitted globally using a 1:1 Langmuir binding model to determine kinetic parameters.

### 3.6 Cell Culture

SupT-1 and HSB-2 cell lines were cultured in RPMI 1640 and IBSS culture media, respectively, each supplemented with 10% FBS and 1% antimycotic-antibiotic. The culturing procedures were performed according to the standard protocol provided by the manufacturer. Cells were cultured in T-75 flasks maintained at 37°C in a humidified atmosphere with 5% CO_2_. The cells were passaged when a flask was 80% confluent and were discarded after 15 passages to avoid senescence-related alterations.

### 3.7 LNP Cytotoxicity *in vitro*

Cytotoxicity of the formulated LNPs was assessed using the CellTiter-Glo assay following the manufacturer’s instructions with slight modifications. Briefly, 100,000 cells were seeded in 96-well plates and incubated overnight at 37 °C in a humidified atmosphere containing 5% CO_2_. The next day, LNP formulations were added at doses ranging from 600 ng to 75 ng per well, and cells were incubated for 24 hours under the same conditions. Following incubation, 100 µL of CellTiter-Glo reagent was added to each well, mixed for 2 minutes on an orbital shaker, and then incubated for an additional 10 minutes to allow stabilization of luminescence signals. Luminescence was recorded to determine cellular ATP levels, and cell viability was calculated as the percentage of ATP relative to untreated control wells.

### 3.8 *In Vitro* Transfection of CD4+ and CD4-T cells

LNPs encapsulating enhanced green fluorescent protein (eGFP) mRNA were evaluated for transfection efficiency in SUPT-1 and HSB-2 cell lines at a standardized dose of 500 ng per 100,000 cells. To minimize adsorptive losses of mRNA LNPs, ssDNA was pre-coated onto the wells of a 96-well plate prior to LNP addition. The prepared LNP formulations were subsequently introduced into the same wells at the specified dose and incubated for 24 hours at 37°C in a humidified atmosphere with 5% CO_2_. Following incubation, cells were harvested into microcentrifuge tubes and washed with HBSS to remove residual culture media and non-internalized LNPs. After washing, the cells were resuspended in HBSS, and the fluorescence intensity, indicative of eGFP expression, was quantified using a fluorescence plate reader.

For the formulation yielding the highest transfection, flow cytometry was performed to quantify both the percentage of transfected cells and the intensity of eGFP expression in SUPT-1 and HSB-2 cells. LNPs surface modified with a non-targeting dT (PolyT) aptamer were included as a negative control. After 24 hours of incubation, cells were collected, washed with HBSS, and stained with a fixable Live/Dead viability dye and anti-CD3 antibody to confirm T cell identity. Samples were analyzed by flow cytometry, and live CD3^+^ cells were gated for eGFP fluorescence. Data were plotted as count versus eGFP intensity on a logarithmic scale, enabling simultaneous evaluation of the percentage of eGFP-positive cells and the distribution of expression levels within each population.

Additionally, the newly designed aptamer sequences generated through the AptaBLE platform were evaluated using the same transfection workflow.

### 3.9 *In vivo* biodistribution

All animal procedures were approved by the North Carolina State University Institutional Animal Care and Use Committee and conducted in accordance with established ethical guidelines. Female C57BL/6 mice, aged 6 to 8 weeks, were maintained on a 12-hour light and dark cycle with free access to food and water. Various formulations of LNPs encapsulating firefly luciferase mRNA were administered intravenously via tail vein injection at a dose of 0.5 mg/kg. Six hours after injection, D-Luciferin was administered intraperitoneally at a concentration of 150mg/kg and allowed to circulate for 10 minutes to enable substrate interaction with the expressed luciferase. Mice were then euthanized, and major organs, including lungs, heart, liver, spleen, kidneys, intestines, femur, and tibia were harvested. Organs were imaged using an IVIS system to detect bioluminescence. Luminescence intensity was quantified for each organ and normalized to signal from saline-treated control animals to evaluate the biodistribution of the LNP formulations.

### 3.10 *In vivo* safety

The toxicity of LNP formulations was assessed by comparing spleen-to-body weight and liver-to-body weight ratios with those of saline-treated control animals. In addition, serum levels of pro-inflammatory cytokines IL-6 and TNF-alpha, as well as the liver injury marker alanine aminotransferase (ALT), were quantified using commercially available ELISA kits according to the manufacturers’ protocols. Briefly, whole blood was collected and allowed to clot at room temperature for 30 minutes, followed by centrifugation at 1200 × g for 5 minutes. The resulting serum was collected, diluted according to the assay instructions, and used for ELISA to determine cytokine and ALT concentrations.

### 3.11 AptaBLE designed CD4 targeting aptamers

AptaBLE is a large language model for predicting aptamer-protein interaction.^41^ Using AptaBLE-MCTS, an algorithm for iteratively exploring and evaluating the broad aptamer search space, a library of 100 40N aptamers predicted to bind hCD4 was produced. The input hCD4 sequence was the recombinant human CD4 (rCD4; Sino Biological, Inc) sequence. To produce a shortlist of aptamers, the secondary structures of all 100 generated aptamers were predicted and clustered using mFold. Tertiary structure analyses of these aptamers in complex with hCD4 was done with Chai-1. Sequences were pairwise-aligned using EMBOSS-Needle. Multiple sequence alignments were done using Clustal Omega.

## 4 Results

### 4.1 Controlled Surface Functionalization of Aptamers on Lipid Nanoparticles

Figure 1A illustrates the schematic process for LNP synthesis. Non-targeted lipid nanoparticles were first generated using a microfluidic mixing device to encapsulate mRNA, followed by surface conjugation with thiolated aptamers. The size and polydispersity index (PDI) of the resulting nanoparticles were measured using DLS (Figure 1B). Non-targeted SM102 and MC3 LNPs had average sizes of 90 and 98 nm, respectively. Aptamer-conjugated formulations (Apt62:25 and Apt62:100) were designed to theoretically attach 25 and 100 aptamers per LNP. However, aptamer conjugation did not significantly alter hydrodynamic diameter, with only a modest size increase of ∼4–12 nm. PDI values for all formulations remained below 0.2, indicating monodisperse particle populations. Figure 1C shows the molecular beacon hybridization assay used to quantify aptamer density on the LNP surface. Aptamer-conjugated LNPs were hybridized with fluorescently labeled molecular beacons, generating a fluorescence signal detected via qPCR. Representative thermal melt curves in Figure 1D show the calibration series of apt62 (0–800 nM). The linear range was identified as 40-47 °C for quantification. Signals from SM102 Apt62:25 and SM102 Apt62:100 formulations confirmed successful aptamer incorporation and enabled quantification of ligand density. The stronger signal observed for SM102 Apt62:100 compared to SM102 Apt62:25 indicates a higher number of aptamers attached when more aptamers are included in the conjugation reaction. In addition to size, PDI and aptamer density, nanoparticles were characterized for zeta potential, and encapsulation efficiency. Complete physicochemical profiles for all LNP formulations used in subsequent studies are provided in Table S2. The average zeta potential of nontargeted LNPs was –2.7 mV, and negatively charged aptamer attachment slightly shifted values to more negative with the values between – 3.0 and –3.8 mV, suggesting minimal effect on surface charge. Encapsulation efficiency across all formulations are above 95%, although the encapsulation efficiency of aptamer-functionalized LNPs appeared relatively lower due to intercalation of the RiboGreen dye into the aptamers.

**Figure 1:**
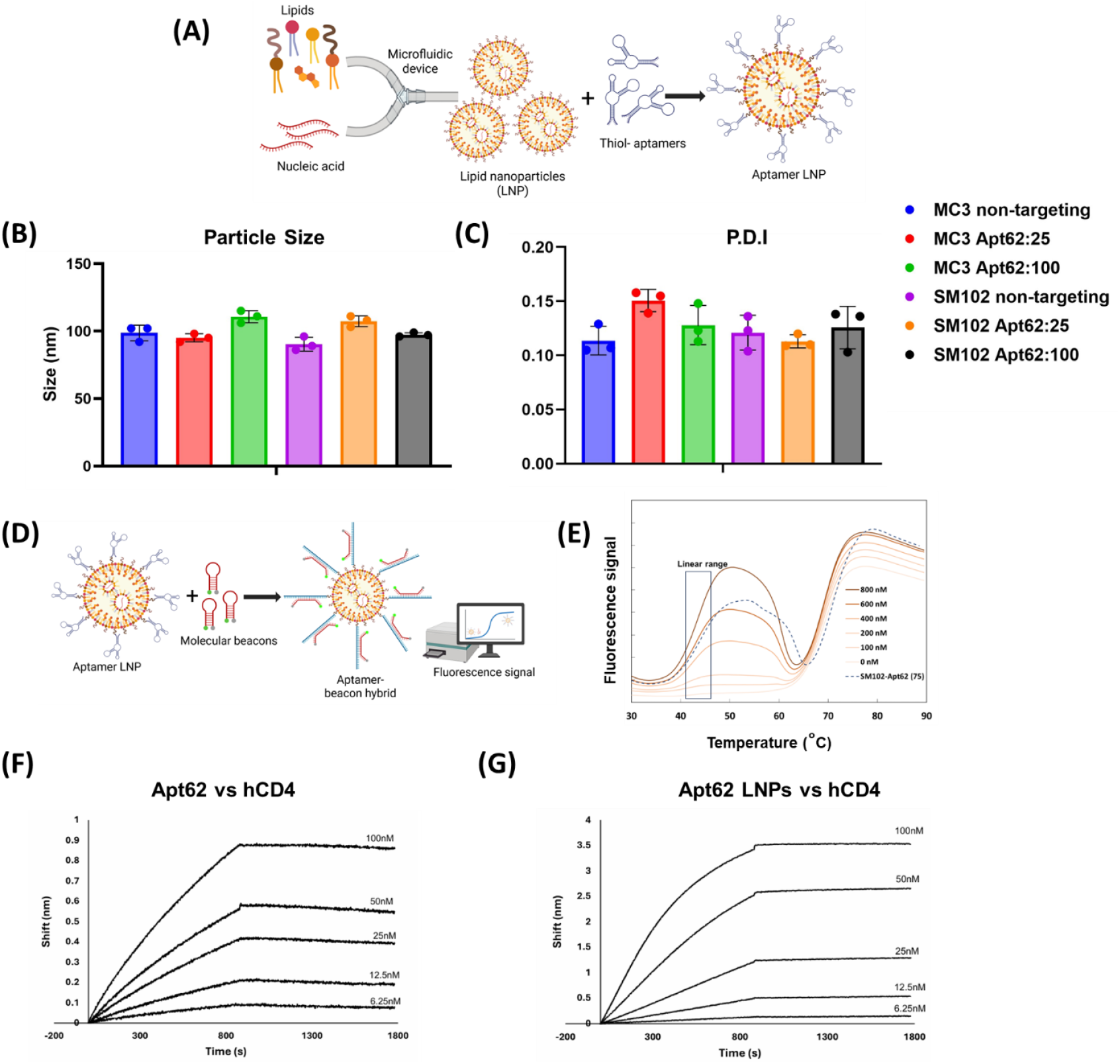
(A) Schematic representation of aptamer–lipid nanoparticle (LNP) formulation using a microfluidic device, followed by thiol–aptamer conjugation. (B) Hydrodynamic size and (C) polydispersity index (PDI) of MC3- and SM102-based LNPs, either non-targeting or Apt62-modified. (D) Schematic illustrating molecular beacon hybridization assay for quantifying aptamer density on LNPs. (E) Thermal melt curves of Apt62 molecular beacons at concentrations ranging from 0–800 nM. (F) Biolayer interferometry (BLI) sensograms of Apt62 binding to recombinant human CD4 protein. (G) BLI sensograms of Apt62-functionalized LNPs binding to recombinant human CD4 protein. Data represent mean ± SD (n=3 for B–C).

In parallel with aptamer-conjugated LNPs, human CD4 (hCD4) and mouse CD4 (mCD4) antibody-functionalized LNPs were synthesized as controls for *in vitro* and *in vivo* studies. Antibody conjugation efficiency was first evaluated by synthesizing eFluor506-labeled hCD4 antibodies conjugated to LNPs during the research and development phase. The conjugation of antibodies was confirmed by quantifying the fluorescence signal of eFluor506-labeled hCD4 antibodies bound to LNPs. Subsequent *in vitro* and *in vivo* experiments were performed using unlabeled antibodies. Attachment of hCD4 mAb to LNPs produced a greater size increase compared to mCD4 mAb. This difference in size may reflect variations in antibody orientation during conjugation, which can influence the apparent hydrodynamic diameter of antibody-functionalized particles. In comparison to antibody-conjugated LNPs, a key advantage of aptamer-conjugated LNPs is the ability to tune surface aptamer density while maintaining physicochemical properties and overall integrity comparable to antibody LNPs.

### 4.2 Aptamer-LNP Conjugation Improves Binding Affinity for hCD4

The binding interaction between Apt62 and recombinant hCD4 was characterized using biolayer interferometry (BLI) binding assay. As shown in **Figure 1F**, Apt62 exhibited concentration-dependent, saturable binding to hCD4, consistent with a 1:1 binding model. The kinetic sensorgrams demonstrated clear association and dissociation phases across the tested concentration range (625 to 20,000 nM), with a calculated equilibrium dissociation constant (Kd) of 536 nM.

To evaluate whether conjugation to LNPs preserved or enhanced binding affinity, Apt62 was immobilized on MC3-based LNPs at a density of 63 aptamers per particle and tested against recombinant hCD4. As shown in **Figure 1G**, the Apt62-LNPs also demonstrated concentration-dependent binding, with markedly stronger signals and slower dissociation relative to the free aptamer. The calculated Kd for Apt62-LNPs was 41.6 nM, representing an approximate 13-fold improvement in apparent affinity. This enhanced binding is attributed to avidity effects arising from multivalent presentation of Apt62 on the LNP surface, which likely enabled the aptamers to overcome steric hindrance caused by the proximity of the binding epitope to the 6xHis tag on CD4.

### 4.3 Aptamer-Functionalized LNPs Demonstrate Safety in T Cells *in Vitro*

Before advancing to functional *in vitro* studies, it was essential to establish a dose range that did not compromise cell viability. To address safety concerns of using aptamer in LNP platform, cytotoxicity was assessed in both CD4^+^ SUPT1 cells which express the target receptor, and CD4^−^ HSB2 cells., which lack CD4 expression. This design ensured that any potential effects of aptamer conjugation on cell survival could be distinguished from receptor dependent uptake or differences in cell lineage. Unconjugated and aptamer-conjugated LNPs containing either MC3 or SM102 were tested across doses from 75 to 600 ng mRNA in LNP per 100,000 cells, using both non-targeted formulations and those functionalized with the highest aptamer density (100 aptamers per LNP). Non-targeting LNPs, lacking aptamers, served as controls. In all conditions, cell viability remained above 80%, indicating that neither LNP lipid composition nor conjugated maximal aptamer display induced significant toxicity in CD4-target positive or CD4-target negative cells (**Figure 2A and 2B)**. A working dose of 500 ng mRNA in LNP per 100,000 cells was selected for all *in vitro* experiments.

**Figure 2:**
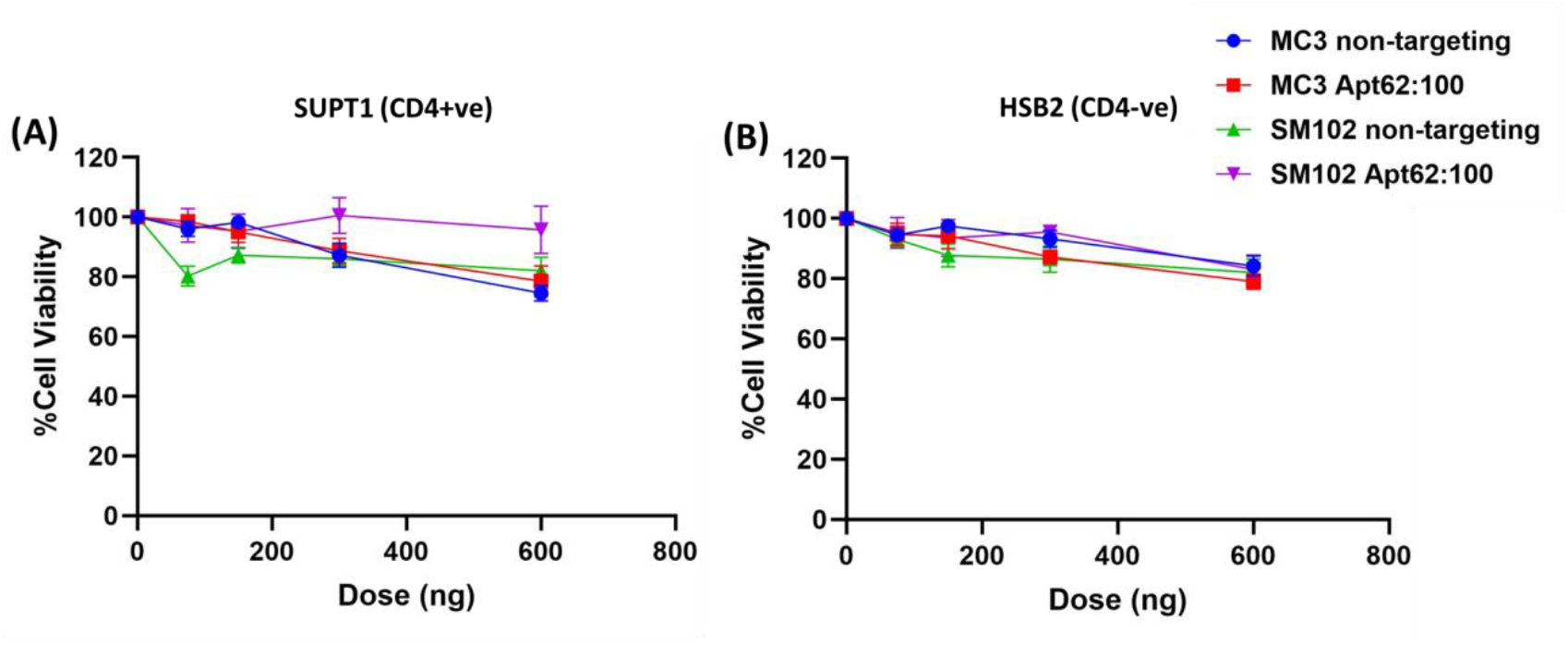
Evaluation of cytotoxicity of aptamer-functionalized lipid nanoparticles (LNPs). Cell viability was measured following treatment with increasing doses of luciferase mRNA-loaded LNPs formulated with either MC3 or SM102 ionizable lipids. (A) SUP-T1 cells and (B) HSB-2 cells were incubated with non-targeting LNPs or Apt62-modified LNPs for 24 hours. Data represent mean ± SD (n=4).

### 4.4 Density-Dependent Aptamer Functionalization Enhances CD4_+_ T Cell Targeting

To evaluate the cell-type selectivity of aptamer-functionalized LNPs, enhanced green fluorescent protein (eGFP) mRNA-loaded nanoparticles were tested in CD4^+^ SUPT-1 and CD4^−^ HSB-2 T-cell lines. LNPs formulated with MC3-based non-targeting LNPs displayed comparable transfection efficiencies in both cell lines, indicating no inherent selectivity **(Figure 3A)**. Conjugation of Apt62 at a 25:1 aptamer-to-LNP ratio significantly increased eGFP expression in SUPT-1 cells, while HSB-2 levels remained unchanged, indicating selective uptake. Increasing the aptamer density to a 100:1 ratio further enhanced SUPT-1 transfection, confirming a density-dependent effect of the aptamer.

**Figure 3:**
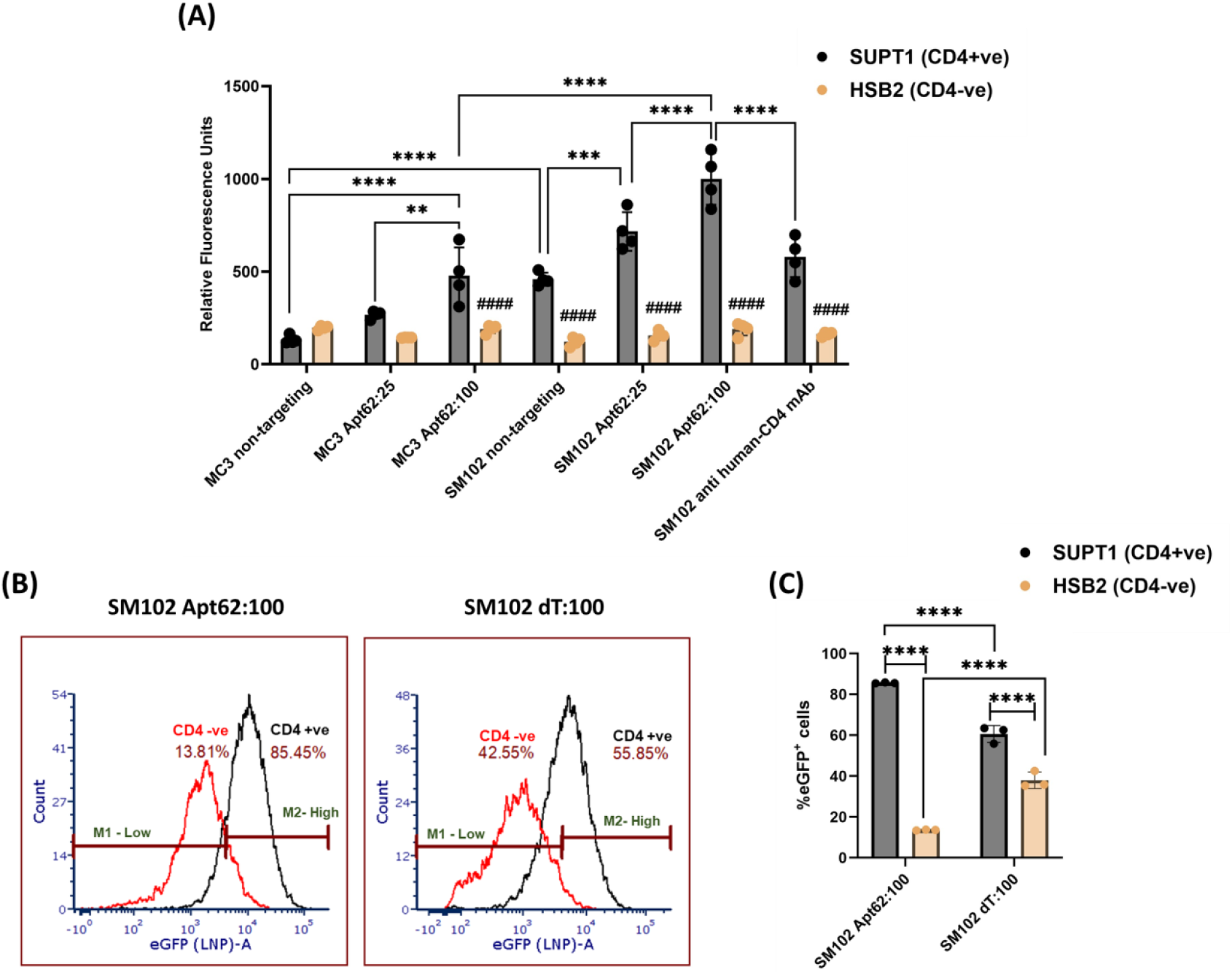
(A) eGFP expression in SUP-T1 (CD4^+^) and HSB2 (CD4^−^) cells following treatment with mRNA-loaded LNPs formulated with MC3 or SM102, either non-targeting, Apt62-modified (25:1 or 100:1), or anti-human CD4 mAb-conjugated. (B) Representative flow cytometry histograms showing eGFP expression in CD4^+^ (black) and CD4^−^ (red) populations treated with SM102 Apt62:100 or SM102 dT:100 LNPs. (C) Quantification of mean fluorescence intensity (MFI) of CD4^+^ and CD4^−^ populations corresponding to panel (B). Data represent mean ± SD (n=4). Statistical significance was determined using two-way ANOVA followed by Tukey’s multiple comparisons test; ****p < 0.0001, ***p < 0.001, **p < 0.01, ####p < 0.0001 versus corresponding CD4^+^ population.

In contrast, SM102 LNPs exhibited elevated baseline transfection in CD4+ SUPT-1 cells even in the absence of targeting ligands, suggesting that the lipid composition itself may contribute to preferential uptake **(Figure 3A)**. Apt62 conjugation further improved CD4+ SUPT-1 targeting in a dose-dependent manner, with the 25:1 and 100:1 formulation showing progressively increased eGFP expression in CD4+ SUPT-1 cells relative to CD4^−^ HSB-2. Notably, SM102 LNPs modified with a CD4 monoclonal antibody, used as a benchmark control, showed lower transfection efficiency in CD4^+^ SUPT-1 cells than Apt62-modified SM102 LNPs.

To confirm cell-type specificity, flow cytometry was performed following transfection with SM102-Apt62:100 and SM102-dT LNPs, the latter serving as a non-targeting control **(Figure 3B)**. Mean fluorescence intensity (MFI) analysis showed that dT-modified LNPs yielded similar eGFP expression levels across both CD4^+^ SUPT-1 and CD4^−^ HSB-2 cells, consistent with non-selective uptake **(Figure 3C)**. In contrast, Apt62-modified LNPs produced a marked increase in eGFP signal in CD4^+^ SUPT-1 cells, with approximately 80% of the population transfected, while CD4^−^ HSB-2 cells remained largely non-fluorescent. These results confirm that Apt62 LNPs enable robust and selective mRNA delivery to CD4^+^ cells.

### 4.5 Aptamer modified LNPs enhance splenic targeting

The biodistribution of luciferase mRNA-loaded LNPs was assessed by quantifying luminescence in major organs following systemic administration. Organs analyzed included liver, spleen, lung, heart, kidney, intestine, femur, and tibia **(Figure 4A)**. Among these, only the liver and spleen exhibited detectable luciferase signals. Non-targeting LNPs formulated with either MC3 or SM102 showed predominant transfection in the liver, consistent with known hepatic uptake of LNPs. No luminescence was observed in the spleen for these non-targeting controls **(Figure 4B)**. Upon conjugation of Apt62, changes in biodistribution were observed. For MC3 LNPs, aptamer modification at a 75:1 ratio resulted in a modest reduction in liver luminescence and a significant increase in spleen signal, with spleen uptake approximately 20-fold higher than non-targeting MC3, suggesting partial redirection away from hepatic clearance toward lymphoid tissue. In contrast, SM102 LNPs maintained high liver signal regardless of aptamer conjugation; however, Apt62-functionalized SM102 LNPs showed a significant increase in spleen-associated luminescence, approximately 80-fold higher than non-targeting SM102, with the Apt62:75 group exhibiting the highest splenic accumulation overall.

**Figure 4:**
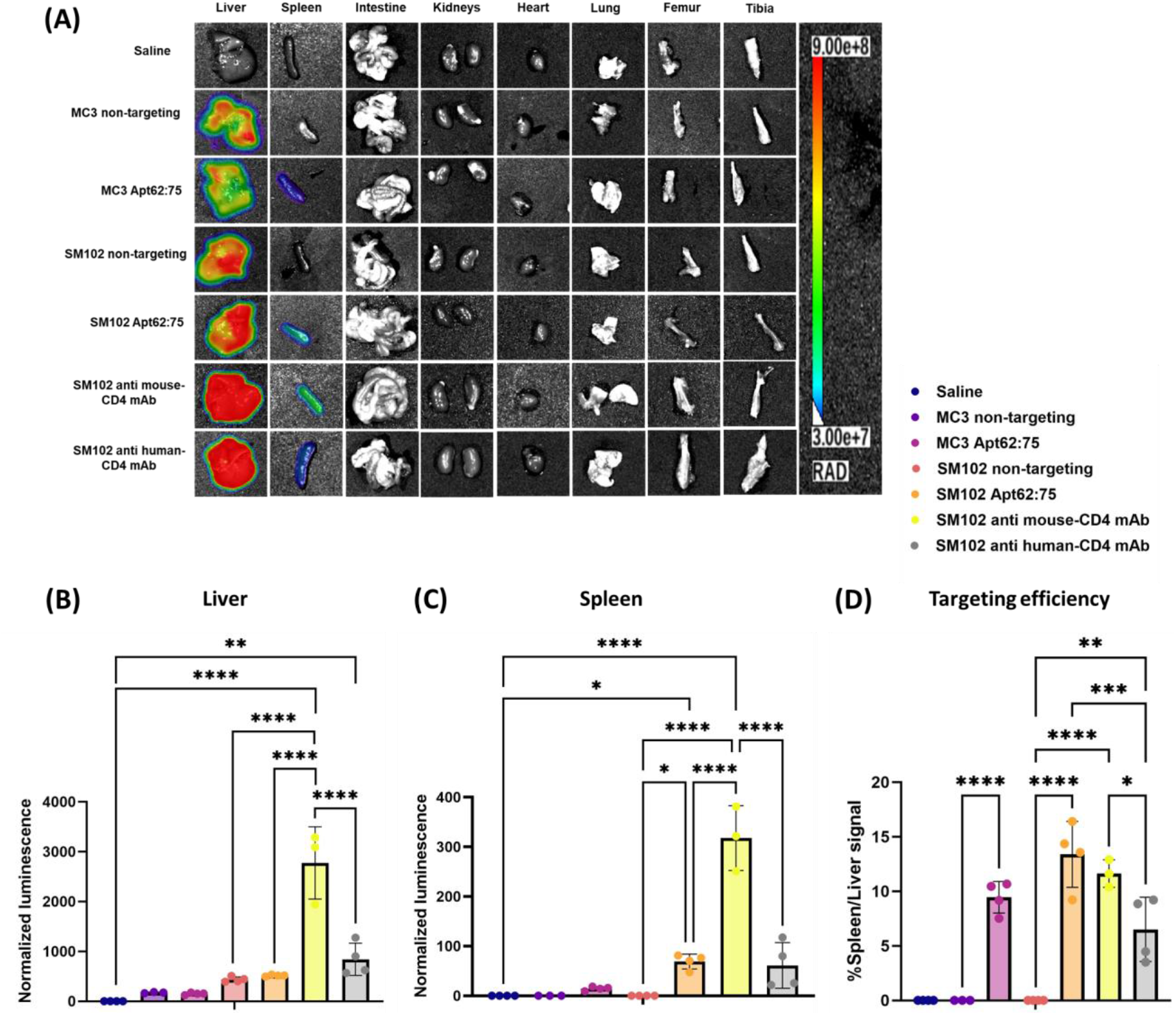
(A) Representative ex vivo IVIS images of major organs collected 6 h post-injection from mice treated with saline, MC3 or SM102 LNPs (non-targeting or Apt62:75), or SM102 anti-mouse/human CD4 mAb-conjugated LNPs. (B) Quantification of normalized luminescence in the liver. (C) Quantification of normalized luminescence in spleen. (D) Percentage of spleen-to-liver luminescence ratio. Data represent mean ± SD (n=4). Statistical significance was determined using one-way ANOVA followed by Tukey’s multiple comparisons test; ****p < 0.0001, ***p < 0.001, **p < 0.01.

To benchmark aptamer targeting against antibody-based approaches, we included two CD4 mAb controls with distinct purposes. Anti–mouse CD4 mAb–LNPs were used to represent conventional antibody targeting in the murine system, while anti–human CD4 mAb–LNPs were evaluated because Apt62 was originally developed against human CD4, allowing comparison between a human-specific aptamer and a human-specific antibody. Anti–mouse CD4 mAb–LNPs increased spleen luminescence by approximately three-to fourfold relative to SM102 Apt62:75 formulation, but this was accompanied by a similar threefold rise in liver luminescence. Because higher spleen signal in isolation can be misleading if accompanied by proportional or greater increases in liver uptake, we quantified targeting efficiency as the spleen-to-liver luminescence ratio. This analysis showed that targeting efficiency **(Figure 4C)** was nearly identical between anti–mouse CD4 mAb–LNPs and SM102:Apt62:75. In contrast, anti–human CD4 mAb–LNPs produced spleen luminescence comparable to Apt62:75 but substantially higher liver signal, resulting in a significantly lower spleen-to-liver ratio. These results indicate that Apt62 conjugation provided stronger splenic selectivity than the human antibody in this murine model, while performing on par with the mouse-specific antibody control.

### 4.6 Aptamer-LNPs Maintain a Favorable Safety Profile, compared to CD4 mAb LNPs

To evaluate the *in vivo* safety profile of the LNP formulations, organ-to-body weight ratios and serum biomarkers were analyzed following systemic administration. No significant differences in liver or spleen weight relative to total body weight were observed across any treatment group, including both non-targeting and Apt62-functionalized LNPs **(Figure 5A and 5B)**, indicating that administration of these LNPs did not induce detectable organ enlargement or tissue burden. Serum cytokine analysis revealed that while IL-6 levels remained largely comparable across most groups, anti-CD4 mAb-conjugted LNPs exhibited a modest upward trend. Importantly, TNF-α was significantly elevated in the anti mouse-CD4 mAb-conjugated LNP group compared to all other formulations, suggesting increased systemic immune activation. Furthermore, ALT levels also showed a modest increase in the antibody-conjugated group, whereas aptamer-modified and non-targeting LNPs remained unchanged relative to saline **(Figure 5C, D, &E)**. Collectively, these results indicate that aptamer-functionalized LNPs maintained a proper safety profile, while antibody-conjugated LNPs were associated with signs of heightened inflammatory and hepatic responses.

**Figure 5:**
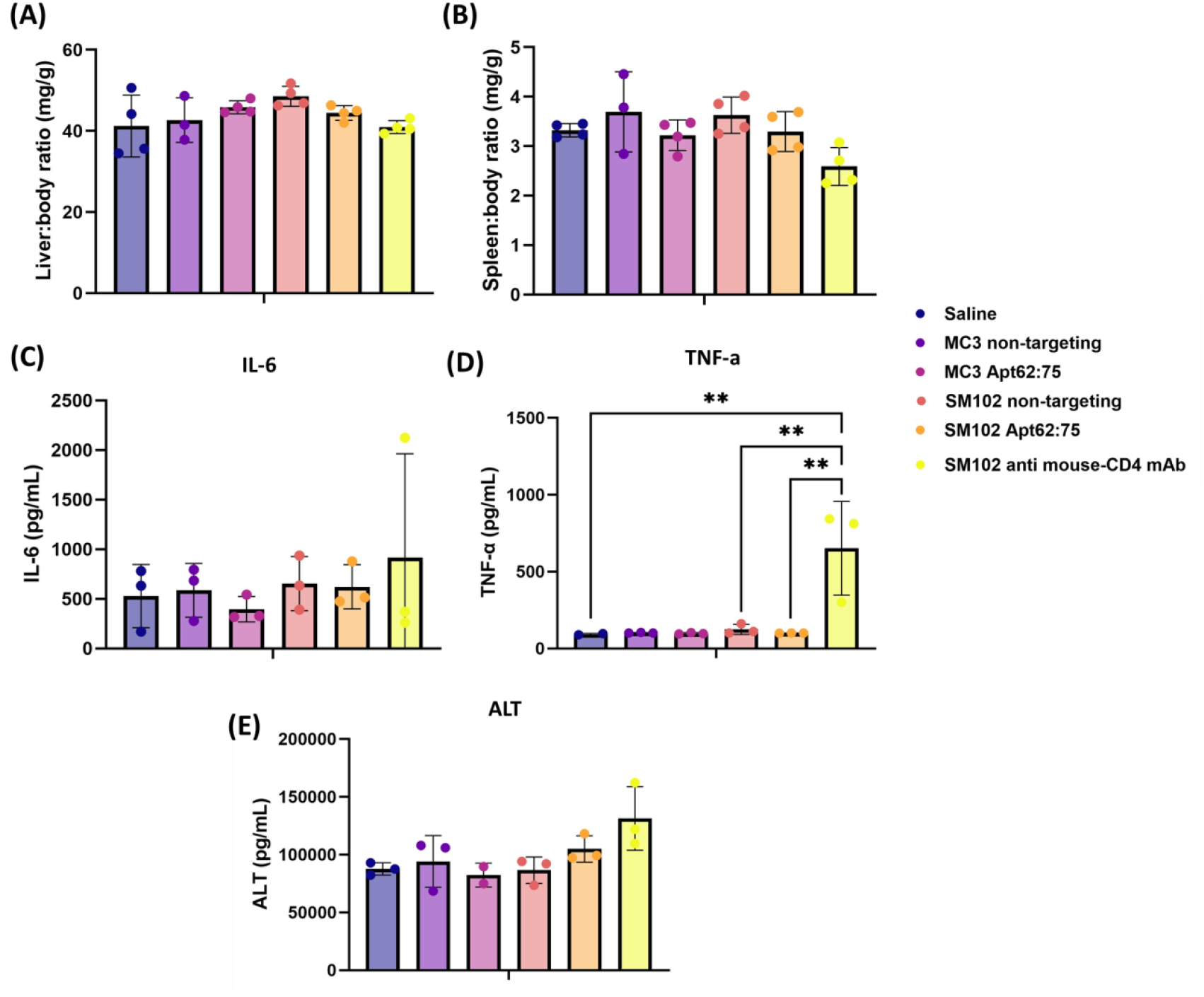
Assessment of organ indices and systemic cytokine markers 6 h post-injection. (A) Liver-to-body weight ratio. (B) Spleen-to-body weight ratio. (C) Serum IL-6 levels. (D) Serum TNF-α levels. (E) Serum ALT levels. Mice were treated with saline, MC3 or SM102 LNPs (non-targeting or Apt62:75), or SM102 anti-mouse CD4 mAb-conjugated LNPs. Data represent mean ± SD (n=4). Statistical significance was determined using one-way ANOVA followed by Tukey’s multiple comparisons test; **p < 0.01.

### 4.7 AptaBLE designed aptamers bind comparably to CD4 as those discovered via *in-vitro* selection

Having established that a previously validated CD4 aptamer reliably confers active targeting to LNPs *in vitro* and *in vivo*, we next asked whether a data-driven design strategy could broaden the ligand repertoire while preserving targeting performance. In this study, we evaluated AptaBLE as a design engine for aptamer ligands enabling cell-specific LNP delivery. Using the AptaBLE-MCTS algorithm, we *in silico* generated a 100-member N40 library (40-nt randomized core) optimized for binding to human CD4 (hCD4) and counter-selected against human albumin to reduce nonspecific serum interactions. AptaBLE-MCTS - a sequence-first language-model search guided by Monte Carlo tree search - efficiently explored the large sequence space while biasing toward high-scoring binders. Library composition was assessed by hierarchical clustering of predicted secondary structures **(Figure 6A)**, revealing broad diversity with only a few tight structural clades; sequence-level similarity was likewise low across candidates (pairwise identity minimal; not shown). Together, these analyses indicate that the designed pool avoids mode collapse and furnishes a structurally and sequence-diverse panel for testing CD4-targeted LNP delivery.

**Figure 6:**
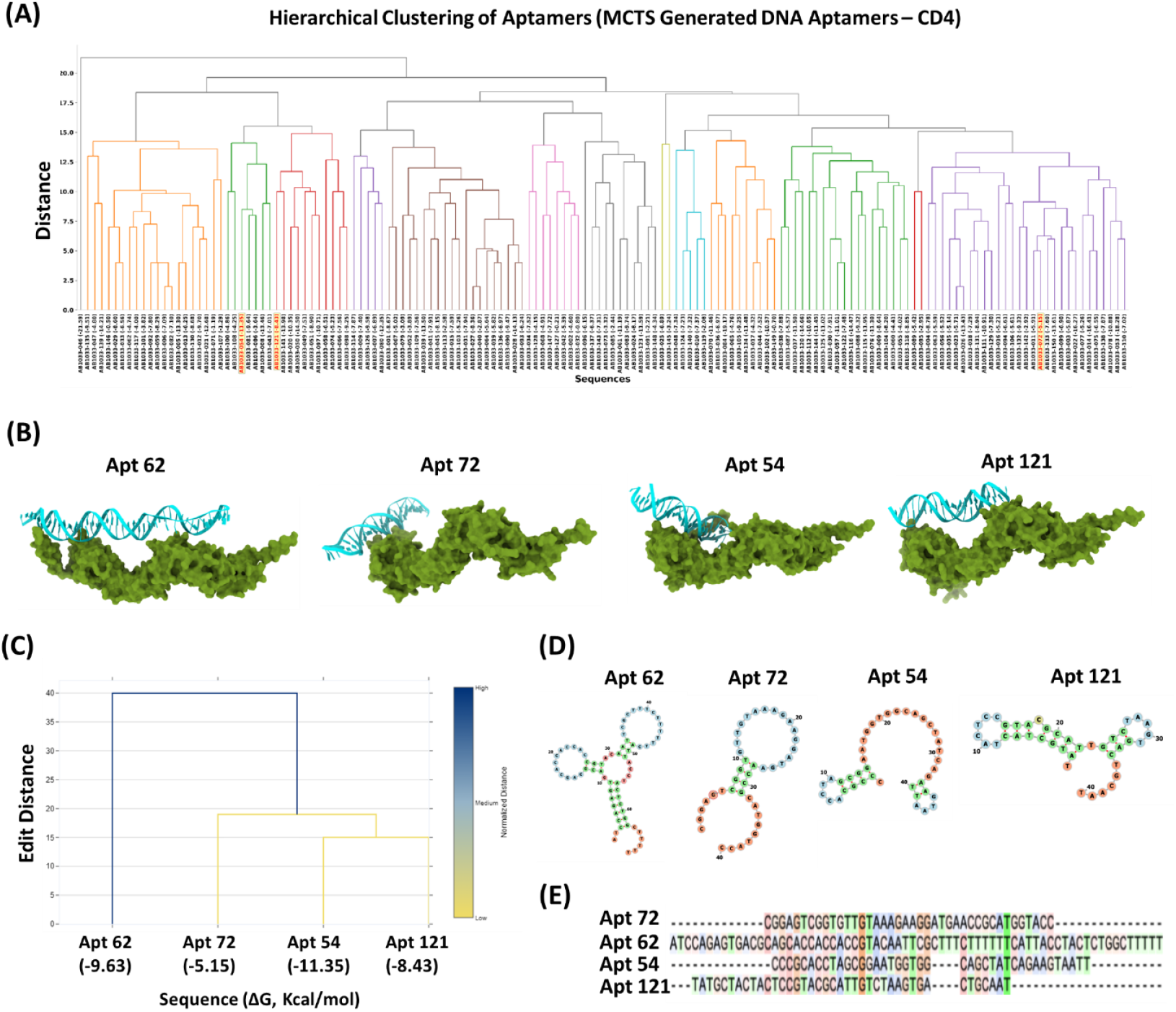
hCD4 aptamer library generated by AptaBLE exhibits high structural diversity and similar predicted binding pose to existing anti-hCD4 aptamer. (A) Dendrogram depicting hierarchical clustering of anti-hCD4 aptamer library predicted secondary structures. Highlighted aptamers were selected and advanced to binding assay. (B) Predicted complexes for hCD4 in complex with 3 AptaBLE-generated aptamers and the lab-selected Apt62. Predicted receptor-binding domains are at the sites of interaction between each aptamer and hCD4. (C) The actual predicted secondary structures (D), and the multiple sequence alignment (E) for all aptamers used in the functional LNP delivery study are shown. For all aptamers used in the functional LNP delivery study. The methodology for the hierarchical clustering is identical to figure 6A. Secondary structure prediction was performed with mFold. Multiple sequence alignment was done with Clustal Omega.

Three novel CD4-binding aptamers were identified for having comparable predicted minimum free energies and GC contents to Aptamer 62. These aptamers were also predicted to bind a similar epitope to Aptamer 62 on the hCD4 surface **(Figure 6B)** in-complex, according to *in-silico* structure docking prediction tools. These candidates were shorter than Aptamer 62 (40 mers) and shared no sequence homology (maximum 40%). Additional analyses show these aptamers share minimal structural similarity or sequential homology to Aptamer 62 **(Figure 6, C-E)**.

Additionally, the three candidate aptamers identified by AptaBLE were conjugated to SM102 LNPs at a 100:1 aptamer-to-LNP ratio and evaluated for selective transfection. These aptamers exhibited similar transfection trends to Aptamer 62, with significantly higher eGFP expression in SUPT-1 cells compared to HSB-2 cells, validating the predictive accuracy of AptaBLE and expanding the functional repertoire of aptamer-guided LNP delivery systems **(Figure 7)**.

**Figure 7:**
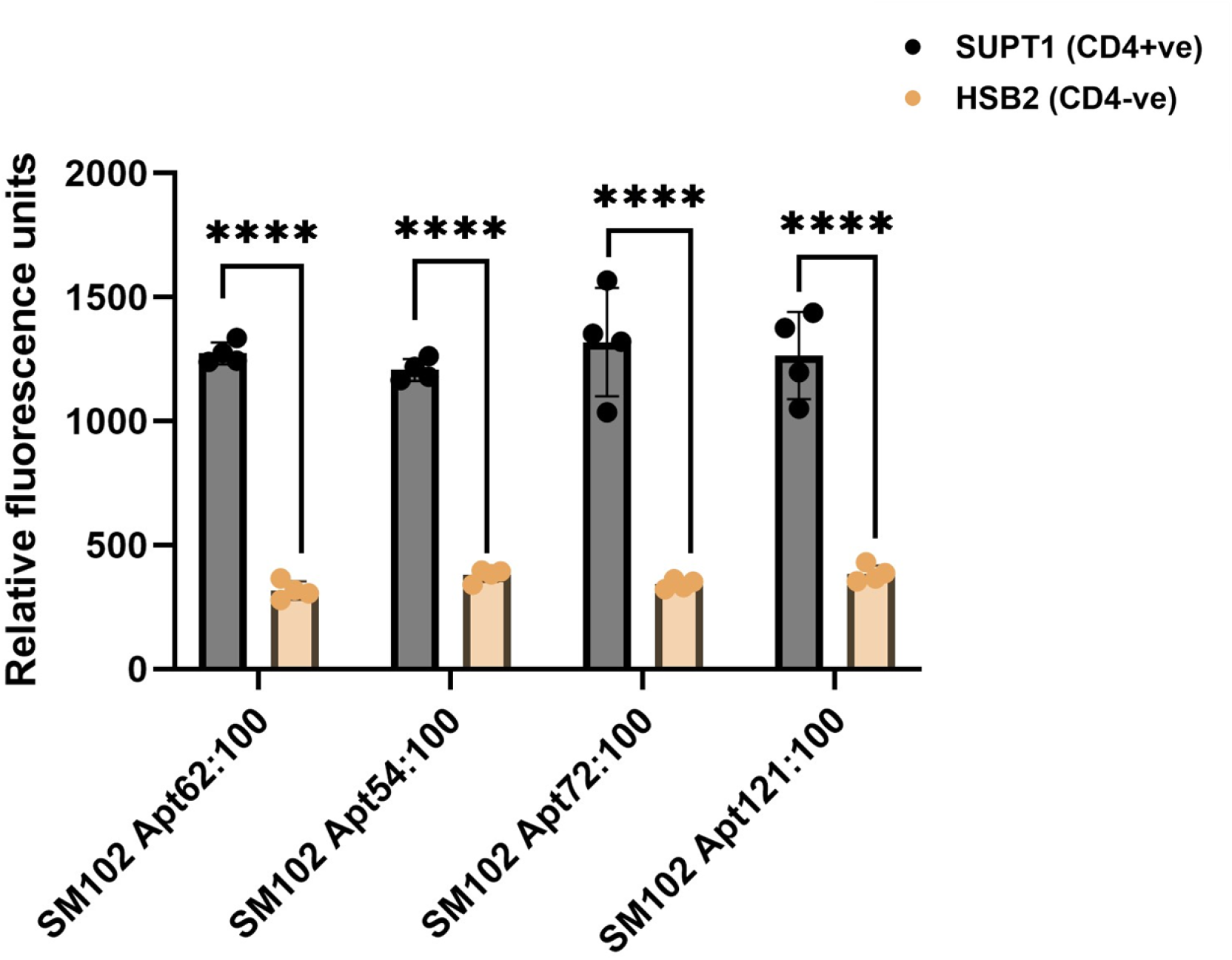
eGFP expression in SUP-T1 (CD4^+^) and HSB2 (CD4^−^) cells following transfection with SM102 LNPs functionalized with AutoNA-predicted aptamers (Apt62, Apt54, Apt72, Apt121) at a 100:1 ratio. Data represent mean ± SD (n=4). Statistical significance was determined using two-way ANOVA followed by Tukey’s multiple comparisons test; ****p < 0.0001.

## 5 Discussion

Active targeting has been widely used for targeted delivery to overcome the challenges of LNP, and multiple ligands have been utilized, such as antibodies, protein fragments, and peptides.^44,45^ Antibodies are attractive for their strong binding affinity and high specificity, but their large size restricts their display density, can destabilize particle structure and packing, complicates purification and scale-up, and increases batch-to-batch variability in manufacturing. In addition, smaller protein fragments, such as Fc regions, can interact with immune cells, altering biodistribution and clearance in ways that compete with receptor-directed uptake.^26,46^ Peptides have also been investigated as targeting ligands, offering smaller size and ease of synthesis, but their utility *in vivo* is often limited by proteolytic degradation, relatively short circulation time, relatively low receptor affinity, along with misfolding and aggregation.^47,48^ These limitations highlight the need for alternative ligands that retain high specificity while being structurally compact, stable in physiological environments, and fully compatible with LNP formulations. In this study, we showed that aptamers can overcome the limitations of antibodies, protein fragments and peptides by redirecting LNPs to deliver mRNA specifically to CD4^+^ cells and immune-rich tissues following systemic administration.

We selected MC3 and SM102, two clinically validated ionizable lipids incorporated into approved or advanced mRNA therapeutics, thereby providing a strong translational foundation.^49,50^ Each formulation contained helper lipid (DSPC), cholesterol, and a PEGylated lipid, with molar ratios adapted from preclinical and clinical studies.^22,50^ After formulating the LNPs, aptamers were conjugated via a maleimide– thiol reaction. The key advantages of aptamer conjugated LNP over antibody conjugated LNP are chemically synthesized aptamer’s tunability and the simplicity of a one-step reaction that maintains uniform orientation. The small size of aptamers and the presence of a single thiol group at the 5′ end enable precise control of the number of ligands attached to each LNP while ensuring correct presentation of the binding domain. We generated LNPs functionalized with 25, 75, or 100 aptamers per LNP, and a molecular beacon complementary to Apt62 confirmed that ligand density increased proportionally with the number of aptamers conjugated. Physicochemical characterization demonstrated only modest increase in particle size and minimal change in zeta potential compared with non-targeting LNPs, indicating that aptamer modification did not alter any physicochemical properties. In parallel, cytotoxicity assays in SUPT1 (CD4^+^) and HSB2 (CD4^−^) cells demonstrated viabilities consistently above 80%, confirming that aptamer display did not compromise cellular health status and that subsequent differences in mRNA delivery would reflect targeting activity rather than nonspecific toxicity.

Receptor engagement is generally the central determinant of selective uptake, and the number of ligands displayed on the nanoparticle surface directly influences delivery efficiency. In SUPT1 (CD4^+^) cells, increasing the density of Apt62 on the LNP surface produced a clear stepwise increase in reporter expression, whereas HSB2 (CD4^−^) cells maintained consistently low levels comparable to non-targeting formulations. This difference between CD4^+^ and CD4^−^ populations confirms that selective delivery arises from ligand–receptor interactions rather than nonspecific internalization. Comparison of MC3 LNP and SM102 LNP formulations further demonstrated that aptamer targeting depends not only on ligand presentation but also on the properties of the underlying ionizable lipid. At equivalent aptamer densities, SM102 LNP consistently produced higher levels of reporter expression than MC3 LNP, a difference likely attributable to its lipid structure, pKa, and membrane fusion characteristics, which together enhance uptake and intracellular trafficking.^50,51^ Benchmarking against antibody-functionalized LNPs highlighted the practical advantage of aptamers. Although anti-CD4 antibody conjugated SM102 LNP showed improvement of mRNA delivery, overall protein expression remained relatively lower than that achieved with SM102 Apt62 LNP, even at reduced aptamer densities. This reduced performance likely reflects the steric bulk of antibodies, which limits the number that can be accommodated per LNP, and their heterogeneous orientation after conjugation, which restricts antigen-binding accessibility. In contrast, aptamers are structurally compact, permit controlled increases in display density, and maintain consistent orientation, allowing more efficient receptor engagement and superior transfection outcomes. Sequence specificity of the aptamer is essential for selective targeting. If the aptamer sequence engages CD4, delivery should be restricted to CD4^+^ cells, whereas a non-binding sequence would be expected to lose this selectivity and distribute more broadly. Consistent with this, SM102 Apt62 LNPs efficiently transfected CD4^+^ SUPT1 cells while maintaining minimal activity in CD4^−^ HSB2 cells, whereas polyT-functionalized LNPs at the same density produced similar levels of reporter activity in both populations. These findings demonstrate that the selective uptake observed with Apt62 is from sequence-specific receptor recognition rather than nonspecific effects of oligonucleotide conjugation.^51^

Our *in vivo* studies demonstrated that aptamer functionalization successfully redirected LNPs to immune-cell–rich tissues. Following systemic administration of MC3 and SM102 non-targeting LNP formulations expressed predominantly in the liver with negligible expression in spleen, tibia, or femur, consistent with ApoE adsorption and LDLR-mediated clearance.^17,52,53^ Apt62 conjugation, however, substantially increased splenic reporter expression compared with non-targeting controls. It is important to consider that the physicochemical properties of the nanoparticles, including particle size and surface charge, remained unchanged between non-targeting and Apt62-functionalized groups; thus, the increased splenic expression can be largely attributed to receptor-mediated uptake. SM102 Apt62 LNPs also showed higher splenic expression than with MC3 Apt62-functionalized LNPs, which is consistent to the performance in CD4^+^ cells *in vitro*. Several groups have developed next-generation lipids, including CICL1 and C14-4, that reduce hepatic sequestration and enable extrahepatic delivery.^44,54^ While our study focused on testing whether aptamer functionalization alone could redirect clinically established, liver-tropic backbones, we anticipate that combining aptamers with these newer lipid scaffolds would provide additive benefits by simultaneously lowering liver expression and enriching delivery to immune-cell–rich tissues.

Benchmarking against antibody-functionalized LNPs demonstrated that aptamers achieve comparable or superior targeting efficiency. SM102 LNPs were functionalized with either mouse or human CD4 antibodies. Both were included for distinct reasons: the mouse antibody provided a species matched control to represent conventional targeting in this model, whereas the human antibody enabled direct comparison to Apt62, which was originally selected against human CD4. Mouse CD4 antibody conjugation produced higher absolute spleen expression than Apt62, but this was accompanied by proportional increases in liver expression, yielding a spleen to liver ratio nearly identical to Apt62 conjugated SM102 LNPs. This demonstrates that greater absolute expression does not equate to improved targeting efficiency when liver remains the dominant site of uptake. The stronger spleen signal with the mouse antibody likely reflects species matched receptor recognition, with Fc dependent interactions further contributing to hepatic clearance. In contrast, the human CD4 antibody produced expression in spleen comparable to Apt62 but disproportionately greater liver accumulation, resulting in significantly less targeting efficiency. The fact that a human selected aptamer achieved targeting efficiency equivalent to a mouse specific antibody, and significantly high to a human antibody, highlights the robustness of cross species recognition. These findings imply that a mouse specific aptamer, or studies conducted in humanized CD4 mice, can further enhance immune cell rich tissue targeting.

Evaluation of systemic safety parameters indicated that aptamer-functionalized LNPs maintained a safety profile comparable to non-targeting LNP formulations. Liver and spleen weights relative to body weight remained unchanged, and both serum ALT activity and cytokine levels were within the same range as saline. These results suggest that incorporation of Apt62 did not induce organ enlargement, hepatocellular stress, or measurable inflammatory activation under the conditions tested. In contrast, antibody-functionalized LNPs showed clear evidence of immune activation. TNF-α concentrations were significantly higher than all other groups, accompanied by modest increases in IL-6 and ALT. This pattern is consistent with systemic inflammatory stimulation together with early hepatic stress. Mechanistically, such effects are likely explained by Fc-dependent interactions, in which antibody Fc regions engage Fcγ receptors on Kupffer cells and splenic macrophages, promoting cytokine release and secondary immune signaling.^25,46,55^

While our data demonstrate that aptamer-functionalized LNPs achieve targeted delivery with a favorable safety profile, these findings should be considered within the broader context of aptamer utility, where historical barriers have constrained clinical translation. As drugs in free form, use of aptamers has been limited by unfavorable PK/PD properties, susceptibility to nuclease degradation, and nonspecific interactions with serum proteins.^56,57^ We addressed several of these barriers by repurposing aptamers as targeting ligands rather than as drugs alone. Beyond the pharmacological limitations of aptamers, a critical barrier lies in their discovery process, where the traditional SELEX workflow remains inherently inefficient.^58^ Iterative bind–wash–amplify cycles extend over weeks to months, enrichment is frequently biased by amplification artifacts rather than true affinity, and insufficient counter-selection often yields polyreactive sequences with broad off-target binding.^58,59^ While refinements such as incorporating negative selection against serum proteins and excluding CpG motifs from candidate libraries can reduce nonspecific interactions and innate immune activation, these strategies do not overcome the fundamental time and resource intensiveness of SELEX.^60–62^

To overcome this discovery bottleneck, we used AptaBLE, our proprietary language model–based platform that can generate de novo high-affinity aptamers within days.^41^ As a proof of concept, four AptaBLE-predicted aptamers were conjugated to SM102 LNPs and tested for their ability to deliver eGFP mRNA selectively to CD4^+^ SUPT1 cells. These aptamer-conjugated LNP formulations showed sequence-specific targeting, producing comparable levels of reporter expression to benchmark Apt62-functionalized LNPs in CD4^+^ cells, while maintaining minimal expression in CD4^−^ HSB2 cells and clearly outperforming non-targeting LNP formulations. These findings confirm that our AI-guided discovery platform can generate novel aptamers that function effectively when displayed on LNPs. This establishes a framework for moving beyond the limitations of SELEX, where in silico–designed aptamers can be rapidly generated and validated in functional contexts, accelerating the integration of aptamer–LNP conjugates into targeted nucleic acid therapeutics.

## 6 Conclusion

This study shows a proof-of-concept of aptamer-functionalized LNPs as a versatile and density-tunable platform for targeted *in vivo* mRNA delivery to CD4^+^ T cells. By functionalizing LNPs with the validated CD4-binding aptamer Apt62, we demonstrated selective binding, enhanced uptake, and increased transfection efficiency in CD4^+^ cells *in vitro*, accompanied by preferential mRNA delivery to immune-rich tissues, particularly the spleen, *in vivo*. Toxicity evaluation confirmed that aptamer-LNPs did not induce measurable hepatic toxicity or systemic inflammatory responses, in contrast to antibody-functionalized LNPs that triggered elevated pro-inflammatory cytokines. Leveraging our proprietary in silico aptamer design framework, AptaBLE, we generated novel, shorter CD4-targeting aptamers that achieved comparable specificity and delivery efficiency to Apt62. These findings highlight the potential of aptamer-functionalized LNPs, augmented by AI-guided aptamer discovery, as a scalable, non-immunogenic, and modular strategy for *in vivo* genetic engineering of T cells, thereby addressing key limitations of current *ex vivo* approaches.

## Acknowledgements

The authors would like to thank Yev Brundo at the University of North Carolina at Chapel Hill for his guidance on *in vitro* and *in vivo* T-cell experimental design, and Isaac Mick along with the staff of the Laboratory Animal Resources Department at North Carolina State University for their assistance with the animal studies.

## Funding

This work was supported by the National Science Foundation SBIR Phase II Grant No. 2127436

## CRediT author statement

**SS:** Conceptualization, Methodology, Validation, Formal analysis, Investigation, Data Curation, Writing original draft, Visualization. **MR:** Methodology, Validation, Formal analysis, Investigation, Data Curation, Writing – Review and Editing. **JD:** Methodology, Validation, Formal analysis, Investigation. **KF:** Investigation, Data curation. **AF:** Conceptualization, Investigation. **YW:** Methodology. **SK:** Investigation. **MB:** Investigation. **SP:** Software, Methodology, Validation, Formal analysis, Investigation. **SY:** Conceptualization, Resources, Writing – Review and Editing, Supervision, Project administration, Funding acquisition.

## Notes

### Competing Interest Statement

The authors have declared no competing interest.

### Summary of Updates

(1) Corrected references and author affiliation (2) Updated figure 6B and Supplementary Table 2

