## Supplementary material for "Lipid Nanoparticles with Aptamers Enable Targeted mRNA Delivery to CD4^+^ T Cells": Supplemenatary information

**Supplementary Table S1: Aptamer sequences**

| Aptamer | Aptamer sequence 5'-3' |
|---|---|
| **Apt62** | ATCCAGAGTGACGCAGCACCACCACCGTACAATTCGCTTTCT TTTTTCATTACCTACTCTGGC TTTTT |
| **Apt 54** | CCCGCACCTAGCGGAATGGTGGCAGCTATCAGAAGTAATT TTTTT |
| **Apt 72** | CGGAGTCGGTGTTGTAAAGAAGGATGAACCGCATGGTACC TTTTT |
| **Apt 121** | TATGCTACTACTCCGTACGCATTGTCTAAGTGACTGCAAT TTTTT |
| **dT (Poly T)** | TTTTTTTTTTTTTTTTTTTTTTTTTTTTTTTTTTTTTTTTTTTTTTTT TTTTTTTTTTTTTTTTTTTTTTTTTT |

**Supplementary Table S2: Physicochemical chemical characterization of Apt-LNPs**

| LNP | Aptamer | mRNA expression | Size (Z-average)/ nm | PDI | Zeta potential/ mV | Encapsulation Efficiency | Ligand density/ nM | Aptamer: LNP ratio* |
|---|---|---|---|---|---|---|---|---|
| SM102 | N/A | fLuc | 86.67 | 0.11 | -2.90 | 97.82 | | |
| | N/A | eGFP | 90.41 | 0.12 | -2.76 | 97.55 | | |
| | Apt62 (25) | eGFP | 94.89 | 0.11 | -3.16 | 98.02 | 285.37 | 38.69 |
| | Apt62 (75) | fLuc | 95.48 | 0.15 | -3.97 | 95.49 | 633.32 | 63.27 |
| | Apt62 (100) | eGFP | 97.15 | 0.13 | -3.74 | 95.32 | 734.00 | 77.01 |
| | dT (100) | eGFP | 97.08 | 0.13 | -4.27 | 95.08 | 579.80 | 64.00 |
| | mouse CD4 (30) | fLuc | 87.14 | 0.13 | -3.91 | 97.71 | | |
| | human CD4 (30) | eGFP | 153.40 | 0.24 | -4.96 | 94.05 | | |
| | Apt54 (100) | eGFP | 99.13 | 0.07 | -4.30 | 95.16 | | |
| | Apt72 (100) | eGFP | 99.70 | 0.09 | -4.30 | 95.88 | | |
| | Apt121 (100) | eGFP | 99.70 | 0.11 | -3.48 | 95.37 | | |
| MC3 | N/A | fLuc | 99.02 | 0.12 | -3.21 | 95.91 | | |
| | N/A | eGFP | 98.65 | 0.11 | -2.74 | 98.03 | | |
| | Apt62 (25) | eGFP | 108.00 | 0.16 | -3.09 | 96.49 | 218.03 | 24.04 |
| | Apt62 (75) | fLuc | 111.40 | 0.15 | -5.39 | 91.60 | 525.63 | 50.30 |
| | Apt62 (100) | eGFP | 110.67 | 0.13 | -5.24 | 94.66 | 565.55 | 66.30 |
